# Proteasome regulation of petite-negativity in fission yeast

**DOI:** 10.1101/2024.05.09.593392

**Authors:** Katie Lin Amberg, Lyrica Hao, Susanne Cranz-Mileva, Mikel Zaratiegui

## Abstract

Mitochondria carry out essential functions in eukaryotic cells. The mitochondrial genome encodes factors critical to support oxidative phosphorylation and mitochondrial protein import necessary for these functions. However, organisms like budding yeast can readily lose their mitochondrial genome, yielding respiration-deficient *petite* mutants. The fission yeast *Schizosaccharomyces pombe* is petite-negative, but some nuclear mutations enable the loss of its mitochondrial genome. Here, we characterize the classical *petite*-positive mutation *ptp1-1* as a loss of function allele of the proteasome 19S regulatory subunit component *mts4/rpn1*, involved in the Ubiquitin-dependent degradation pathway. The mutation results in an altered oxidative stress response, with increased levels of oxidized glutathione, and increased levels of mitochondrial and cytoplasmic chaperones. We propose that Ubiquitin-proteasome regulation of chaperones involved in the Unfolded Protein Response and mitochondrial protein import underlies petite-negativity in fission yeast.

## Introduction

Mitochondria contain the remnants of their ancestral alpha-proteobacterial genome. While much reduced, after shedding the majority of its genetic content by outright loss or transfer to the nuclear genome, it still encodes a number of proteins as well as complete translational ncRNA machinery (rRNA and tRNA) (Gray 2012). Shortly after the description of the mitochondrial genome (mtDNA), it was discovered that the respiration deficient small colonies that arise spontaneously in *Saccharomyces cerevisiae*, known as *Petites*, harbor mutations in or complete loss of mtDNA (Corneo, et al. 1966; Moustacchi and Williamson 1966). The study of *petite* mutations have led to insights into mitochondrial involvement in metabolism, aging, and disease.

*Petite* positivity is the ability for yeast cells to tolerate the loss of their mtDNA. Conversely, *petite* negativity is the inability to tolerate the loss of mtDNA. Testing for *petite* positivity involves multiple rounds of growth in media containing Ethidium Bromide (EtBr) which is highly toxic to mtDNA and will result in the formation of *petite* colonies when plated onto glucose media (Slonimski, et al. 1968; Haffter and Fox 1992). Growth in EtBr results in the formation of mitochondria devoid of mtDNA (ρ0). The exact mechanism of EtBr induced loss of mtDNA is unknown but is suspected to involve specific inhibition of mtDNA replication and transcription (Zylber, et al. 1969). Since multiple essential subunits of the Electron Transport Chain (ETC) complexes are encoded in the mtDNA, these mitochondria are deficient for oxidative phosphorylation, and cells harboring only ρ0 mitochondria are unable to perform respiration. *Petite* positive species are able to survive as ρ0 as long as they are in the presence of Glucose, which provides ATP by way of fermentation, but are unable to survive in respiration carbon sources such as Glycerol or Pyruvate (Slonimski, et al. 1968; Haffter and Fox 1992).

The *S cerevisiae* mtDNA codes for the subunits of the ETC (*cob1*, *cox1*, *cox2* and *cox3*) (Gray 2012; Freel, et al. 2015). These are core membrane-bound subunits that are essential for the function of complex III and IV of the ETC. Loss of mtDNA results in a loss of function of complexes III (*cob1*) and IV (*cox1*/*cox2/cox3*). Since *S cerevisiae* lacks complex I this results in dysfunction of the ETC and prevents the pumping of protons into the intermembrane space of the mitochondria from the matrix. The only ETC component left intact is complex II, succinate dehydrogenase, which transfers electrons to the Quinone pool but does not pump protons. Additionally, the mtDNA genes *atp6*, *atp8* and *atp9* code for essential components of the membrane bound proton pore F_o_ subunit of the F-ATP synthase, resulting in inability of ρ0 cells to utilize the proton gradient to generate ATP. Together, this catastrophic loss of oxidative phosphorylation results in respiration deficiency and a large drop in mitochondrial inner membrane potential ΔΨ (Liu, et al. 2022). Since protein import into the mitochondrial matrix depends on ΔΨ, essential mitochondrial functions, like Iron-Sulfur cluster biosynthesis, are also compromised (Veatch, et al. 2009).

*Petite* mutants are capable of foregoing oxidative phosphorylation in the presence of glucose for fermentation but are somehow capable of maintaining enough ΔΨ to support some measure of mitochondrial protein import (Chen and Clark-Walker 2000). *Petite*-positivity in *S cerevisiae* depends on the action of several nuclear genome encoded factors. Conversely, some *petite*-negative yeasts like *Kluyveromyces lactis* and *Schizosaccharomyces pombe* can be made *petite*-positive by nuclear mutations. The study of *petite*-supressing and *petite*-enabling mutations has revealed that mitochondrial function can be supported by the inverted action of the nuclear-encoded F_1_ subunit of the ATP synthase and the ATP/ADP exchange machinery, which can maintain ΔΨ by hydrolysis of ATP produced by the fermentative pathway (Chen and Clark-Walker 2000). *Petite*-positivity in *S cerevisiae* is lost in mutants in the a and b subunits of F_1_-ATPase and in the *PET9* ATP/ADP translocase (Chen and Clark-Walker 1999, 2000). Similarly, mutation of the *YME1* iAAA protease results in increased levels of Inh1, which inhibits ATP hydrolysis by the F_1_-ATPase subunit, resulting in *petite*-negativity (Thorsness, et al. 1993; Kominsky, et al. 2002). Other mitochondrial chaperones acting in the intermembrane space also are involved: Tim9/Tim10, which shuttle cargo between the outer (TOM) and inner (TIM) membrane protein translocation complexes and are regulated by Yme1 (Spiller, et al. 2015), are also necessary for *petite* positivity in *S cerevisiae* (Senapin, et al. 2003). Several *petite*-enabling mutations in *K lactis* and *S pombe* have also been identified, painting a consistent picture. In *K lactis*, inactivation of INH1 and mutations of the α, β and γ subunits of F_1-_ATPase that increase ATP hydrolysis allow for growth of ρ0 cells(Clark-Walker 2007). In *S pombe*, mutants of the β and γ subunits of F_1_-ATPase also become *petite*-positive(Li, et al. 2019; Dinh and Bonnefoy 2023), but their effect on ATP hydrolysis activity has not been reported. Notably, overexpression of *S cerevisiae* Yme1 in *S pombe* renders it *petite*-positive (Kominsky and Thorsness 2000).

A survey of gene essentiality suppressors in *S pombe* revealed a network of interesting genetic interactions between factors involved in mitochondrial translation and the proteasome (Li, et al. 2019). A mutation in the 19S regulatory subunit component *mts4* (*RPN1* in *S cerevisiae*), as well as overexpression of Mts4 and several other proteasome related factors, enabled growth of ρ0 cells after EtBr treatment. This indicates that the proteasome activity can regulate survival in the absence of mtDNA in fission yeast. However, the mechanism of this phenomenon was not addressed. Mutations of proteasome subunits in *S cerevisiae* also affect either the stability of the mtDNA or the survival of ρ0 cells: loss of *RPN4*, a transcription factor that drives expression of proteasome factors, increases survival to EtBr treatment (Dunn, et al. 2006), and deletion of the proteasome activator *BLM10* and the proteasome chaperone *UMP1* increase the frequency of *petites* (Sadre-Bazzaz, et al. 2010). The 26S proteasome, composed of the 19S regulatory subunit and the 20S core subunit, affects multiple cellular pathways by regulating the levels of cellular factors through the Ubiquitin-proteasome degradation pathway (Finley, et al. 2012). The 20S core proteasome can also degrade misfolded and oxidized proteins independently of the 19S subunit (Raynes, et al. 2016). Since the loss of ΔΨ in ρ0 cells results in cytoplasmic accumulation of mitochondrial protein precursors and the activation of the Unfolded Protein Response (UPR) (Narayana Rao, et al. 2022), 20S proteasome activity may be important for their survival. However, the wide range of substrates of the proteasome makes it difficult to propose a mechanism for its involvement in *petite*-positivity.

The *ptp1-1* mutation was the first nuclear *petite*-enabling mutation described in *S pombe* (Haffter and Fox 1992). It was isolated as a background enabling a high frequency of ρ0 cells after incubation in EtBr. It has since been used to investigate mitochondrial genetics in multiple studies (Dinh and Bonnefoy 2023), but has remained unidentified. In order to investigate the mechanism of *petite*-negativity in fission yeast, and to further inform the phenotypes of ρ0 cells obtained with this mutation in the presence of other mutations of interest, we set out to identify and characterize this mutation.

## Materials and methods

### Testing for *petite*-positivity

To assay strains for *petite* positivity, 5ml cultures of YES supplemented with 12.5 ug/ml EtBr and 2% potassium acetate (Haffter and Fox 1992) are inoculated with ρ+ cells and left to incubate at 30°C until saturated (up to 2 days). 150μl of saturated culture are used to inoculate 5ml of the same media and incubate for up to 5 more days (total 1 week in EtBr media). The cells were centrifuged, resuspended in 200μl of media and plated onto YES with 3% glucose and incubated at 32°C. Large colonies (ρ+ which have survived EtBr treatment) appear within a few days. Smaller *petite* colonies that appear after about 1-2 weeks of incubation were restreaked in fermentation (YES with 3% glucose) and in respiration (YES with 3% glycerol and 0.1% Glucose) media. *Petite* respiratory negative colonies were confirmed as ρ0 by colony PCR with primers amplifying the *atp6* gene.

### Strains and genetic crosses

All strains used in this study are detailed in table 1. To characterize the ptp1-1 mutation, the PHP14 strain (Haffter and Fox 1992) was first crossed with a ρ+ strain, and the segregants obtained by random spore analysis were subjected to the EtBr treatment described above to identify ptp1-1 ρ0 strains. The cross was repeated twice more. Asci from the final cross was dissected to obtain full tetrads, and the segregants were phenotyped for *petite* positivity. Two full tetrads showing 2:2 segregation of *petite* positivity were chosen for full genome resequencing.

**Table 1.** Strains used in this study.

| strain | mating type | genotype | Source |
| --- | --- | --- | --- |
| PHP14 | h- | *ρ0, ade6-M216, leu1-32, ptp1-1* | Dr. Thomas Fox |
| FWP16 | h+ | *ura4-D18* | Dr. Charles Hoffman |
| ZB4660 | h+ | *ura4-D18, mts4L636P* |  |
| ZB4414 | h+ | *mts4L636P* |  |
| ZB4419 | h+ | *mts4L636P* |  |
| 975 | h+ |  | Dr. Nancy Walworth |
| ZB4622 | h+ | *atg43-1::kanMX6, sdh2-mCherry::KanMX, cpy1-venus::natMX, CFP-atg8::leu1+, his3-D1?* |  |
| ZB4624 | h+ | *ptp1-1, atg43-1::kanMX6, sdh2-mCherry::KanMX, cpy1-venus::natMX, CFP-atg8::leu1+, his3-D1?* |  |

### DNA purification

Cells were grown to saturation in rich media, and 50OD (approximately 5e8 cells) was used for genomic DNA extraction with the NEB monarch gDNA extraction kit, with the following modifications: The cells were first washed twice in water, and then resuspended in CSE buffer (150mM Citrate buffer pH5.4, 1.2M Sorbitol, 10mM EDTA) with 0.4mg/ml Zymolyase 100T, and incubated at 37C for 45 minutes to obtain spheroplasts. Spheroplasts were pelleted and resuspended in 500 ul of 5xTE, lysed by the addition of 10% SDS to a final concentration of 1%, and incubated at 50C for 1h. The lysates were cooled in ice, precipitated by addition of 175ul of 5M Potassium Acetate, and further incubated in ice for 15 minutes. After centrifugation at 18000g, the supernatant was pipetted to fresh tubes through a piece of miracloth, and nucleic acids were precipitated by addition of 500ul cold isopropanol, incubation at −20C for 30m and centrifugation for 30m at 18000g. The pellets were washed in 70% Ethanol, resuspended in 200ul 5xTE with 100ug/ml RNAseA and incubated at 37C for 30m. Then 6ul of 10% SDS and 40ug ProteinaseK were added, followed by 50C incubation for 60m. After that, the extract was mixed with 400ul NEB monarch kit binding buffer and loaded onto a Monarch gDNA purification column. DNA purification from this point on followed manufacturer’s recommendations.

### High Throughput Sequencing

Approximately between 1 and 4 ug of genomic DNA was first sheared to a target size of around 150bp in a microTUBE AFA fiber pre-slit snap-cap Covaris tube in the Covaris M220 focused ultrasonicator (150W Peak Incident Power, 10% duty Cycle, 200 cycles per burst, 430s total time). The DNA was then purified with 1.8 volumes of MagBio HighPrep magnetic beads. 1ug of sheared DNA were used to prepare barcoded libraries using the NEB ultra II DNA library prep kit, pooled and sequenced in a MiSeq 2×150 v2 flow cell.

### Sequence analysis

The fastq files were analyzed for polymorphisms using the CrossBow 1.2.1 pipeline (Langmead, et al. 2009) using the *Schizosaccharomyces pombe* ASM294v2 assembly as a reference. The SOAPsnp output generated by Crossbow was analyzed for polymorphisms present in the *petite*-positive segregants and absent in the *petite*-negative segregants.

### Plasmid generation

Oligonucleotides are specified in Table 2. The Crispr plasmid for editing the *ptp1-1* mutation was created via PCR using anti WT primers (oM2631/oM2632) to insert the gRNA targeting sequence site into crispr plasmid pMZ374 (adh1Cas9-pgRNA *ura4*). Similarly, the crispr plasmid for reverting from *ptp1-1* back to WT was created via PCR using anti ptp1 primers (2633/2634) to insert into plasmid pMZ377 (adh1Cas9-pgRNA LEU2).

**Table 2.**
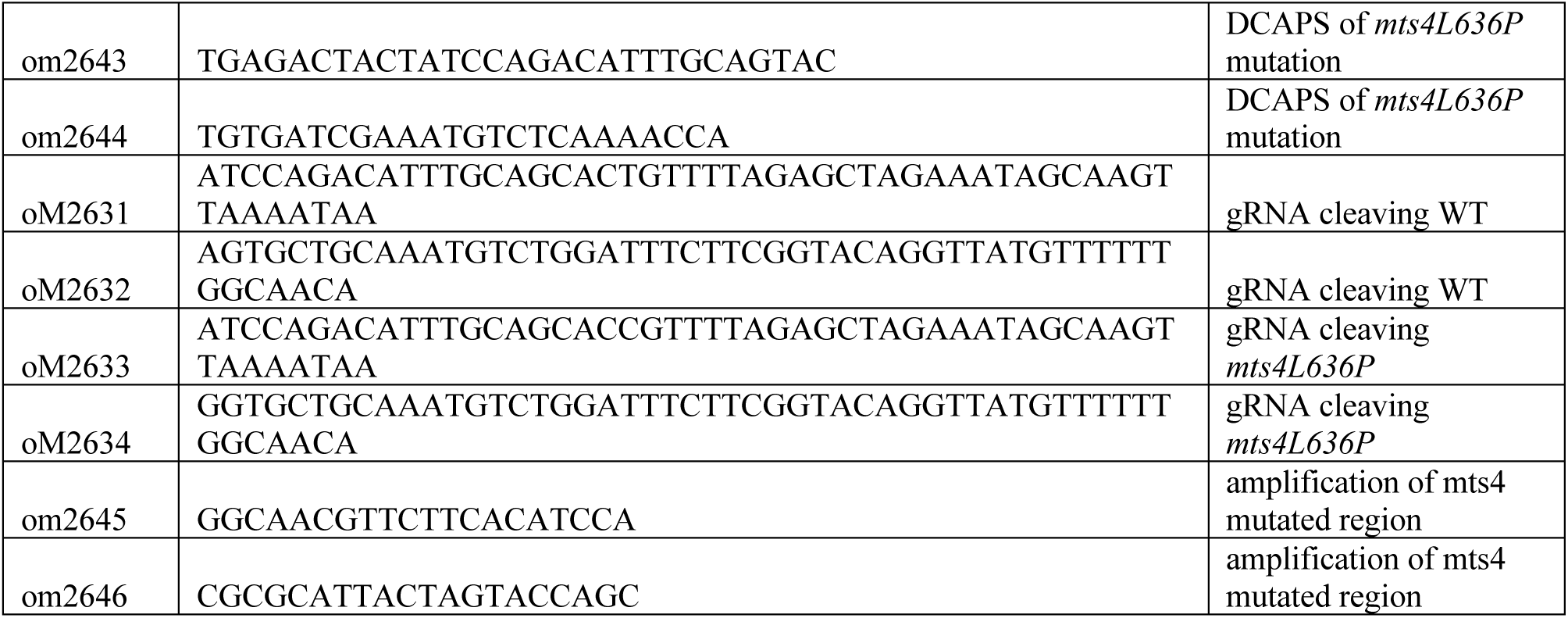
Oligonucleotides.

### CRISPR mutagenesis

CRISPR mutagenesis was performed as previously described (Jacobs, et al. 2014). Plasmid transformation was carried out with the Lithium Acetate method using 600ng of plasmid (containing Cas9 and the gRNA expression cassette) and 400ng of Homologous Recombination donor. The cells were plated onto selection media and grown for a week at 32C. Small colonies (representing Cas9-expressing survivors) were restreaked on selective media and genotyped by DCAPS PCR (oligonucleotides oM2643/2644 followed by digestion with ScaI). The target region was amplified (oM2645/2646) from mutant candidates and subjected to Sanger sequencing to confirm the intended mutations.

### G0 survival

To analyze the survival on G0, prototrophic WT and *ptp1-1* mutants cells grown in 5ml of EMM media with 3% Glucose to a cell density of 5e6 cells/ml, harvested by centrifugation and washed 3 times in G0 media (EMM media without Ammonium Chloride and 3% Glucose), then resuspended in the original volume of culture in G0 media, and returned to shaking at 32C. Samples were taken before and at 48h intervals after the G0 media switch, seeded at 1000 cells per plate in YE agar plates, grown for 2 days at 32C, and counted for colony forming units (cfu). At certain time points, a 5ul sample of the cultures was spread as the inoculum in YE plates, and cells were individually dissected from the inoculum and arranged in a 9×9 grid using an MSM400 tetrad dissector.

### Proteasome assay

To directly assay proteasome activity, cultures from WT or *ptp1-1* mutants were grown in YES or YES/2% KAc/EtBr media to 1e7 cells/ml density, and 4e8 cells were harvested by centrifugation, washed once in ice-cold water, and resuspended in 500ul extraction buffer (50mM Tris pH 7.5, 5mM MgCl_2_, 1mM DTT, 1/100 dilution of protease inhibitor cocktail IV (Millipore)) with or without 1mM ATP. The cell suspension was vortexed with 500ul of 0.5mm Zirconia beads for 10 cycles of 1m vortexing at 3000rpm and 1m cooling on ice. The bead/lysate mixture was centrifuged at 14000 rpm at 4C for 15 minutes, and 300ul of clarified lysate was recovered, avoiding the precipitated lipid layer. The lysate was further clarified by centrifugation at 14000rpm at 4C for 10 minutes. Protein concentration was determined by Bradford assay along with a BSA standard, and the concentrations were normalized by addition of an appropriate volume of extraction buffer to the most concentrated samples.

The proteasome assay was carried out by mixing 50ug of extract at 1ug/ul concentration with 150ul of assay buffer (final concentrations 50mM Tris pH 7.5, 5mM MgCl_2_, 1 mM DTT, 50uM Suc-LLVY-AMC substrate) with or without 1mM ATP. Control reactions in the presence of 50uM MG132 (a proteasome inhibitor) were carried out in parallel. The reactions were performed in white 96 well plates in a Biotek Synergy HT plate reader at 32C. Fluorescence was monitored with excitation at 360nm and emission at 460nm, every 2 minutes for 2 hours. Proteasome activity was determined by calculating the rate of MG132 sensitive fluorescence release by linear regression over time. We performed 3 biological replicates of each strain or condition, repeated over 3 technical replicates.

For native gel proteasome activity assay, 50ug of extract was subjected to electrophoresis in 3.5% polyacrylamide gels as previously described (Roelofs, et al. 2018), developed in 50mM Tris pH7.5, 5mM MgCl2, 1mM ATP and 50uM Suc-LLVY-AMC with or without 0.02% SDS, and imaged in a Kodak GelLogic 1500 system with a 530DF100 filter.

### Proteomics analysis

Three biological replicates each of WT and *ptp1-1* mutants were grown in YES media with 2% Potassium Acetate and 12.5ug/ml EtBr to a cell density of 1e7 cells/ml. Then, Trichloroacetic Acid (TCA) was added from a 50% w/v solution in water to 2e8 cells of culture to a final concentration of 20% TCA, harvested by centrifugation and washed once in 20% TCA. Pellets were resuspended in 20% TCA, combined with 0.5mm Zirconia beads and vortexed for 10 rounds of 1 minute vortexing at 3000rpm followed by 1 minute cooling on ice. The lysates were recovered from the beads by piercing the bottom of the tube and spinning at 1500g in a swing-bucket centrifuge, and the beads were washed with 500ul of 20%TCA, combining the flowthrough. The precipitates from the lysate were recovered by centrifugation at 20000g for 15 minutes. The pellet was washed twice in 100% acetone, then dried and sent for whole proteome analysis by SDS-PAGE separation, protease digestion of gel slices, and Liquid chromatography/Mass Spectrometry (LC/MS). Statistical analysis of LC/MS peptide abundance data was reported as log2 ratios and significance calls (p values) without and with multiple comparison corrections (Bonferroni, Holm-Sidak, Benjamini-Hochberg FDR, and Strimmer’s LFDR)

### Metabolomics

For Metabolomics analysis, an aliquot of the same cultures harvested for proteomics were processed as follows: 5e7 cells were harvested by filtration onto 0.45um pore, 47mm diameter Nylon filters. The filters were then overlaid, cells face down, onto 1.2ml of an ice-cold 40:40:20 mixture of Acetonitrile:Methanol:H_2_O_2_ in a 5cm petri dish sitting on crushed ice. The mixture was neutralized by addition of 83ul of 15% NH_4_HCO_3_, and incubated for 15 minutes on ice. The cells were washed onto the extraction buffer by pipetting over the surface of the filter, and then recovered from the petri dish into an eppendorf tube. The extracts were then centrifuged for 15m at 20000g and 4C, and 700ul of the clarified supernatant were recovered and sent for positive and negative polarization LC/MS analysis.

## Results

### *ptp1-1* is a loss of function mutation in *mts4*

After two rounds of backcrosses of the original PHP14 *ptp1-1* isolate, following the *petite*-positive phenotype of the segregants to clean the background of possible mutations caused by the original EtBr treatment, we isolated two complete tetrads that showed a 2:2 segregation of *petite* positivity. We sequenced the genome of all 8 segregants and analyzed the polymorphisms present. Only one polymorphism was present in all 4 *petite*-positive segregants and absent in all 4 *petite*-negative segregants. This polymorphism results in a mis-sense mutation in the protein coding gene *mts4*, changing Leucine 636 to a Proline (Figure 1A, B, C). Of note, a different mutation in *mts4* has been previously described to confer *petite*-positivity in fission yeast (Li, et al. 2019) (Figure 1B). Therefore, the *mts4L636P* mutation is a candidate for the identity of the *ptp1-1* allele.

**Figure 1.**
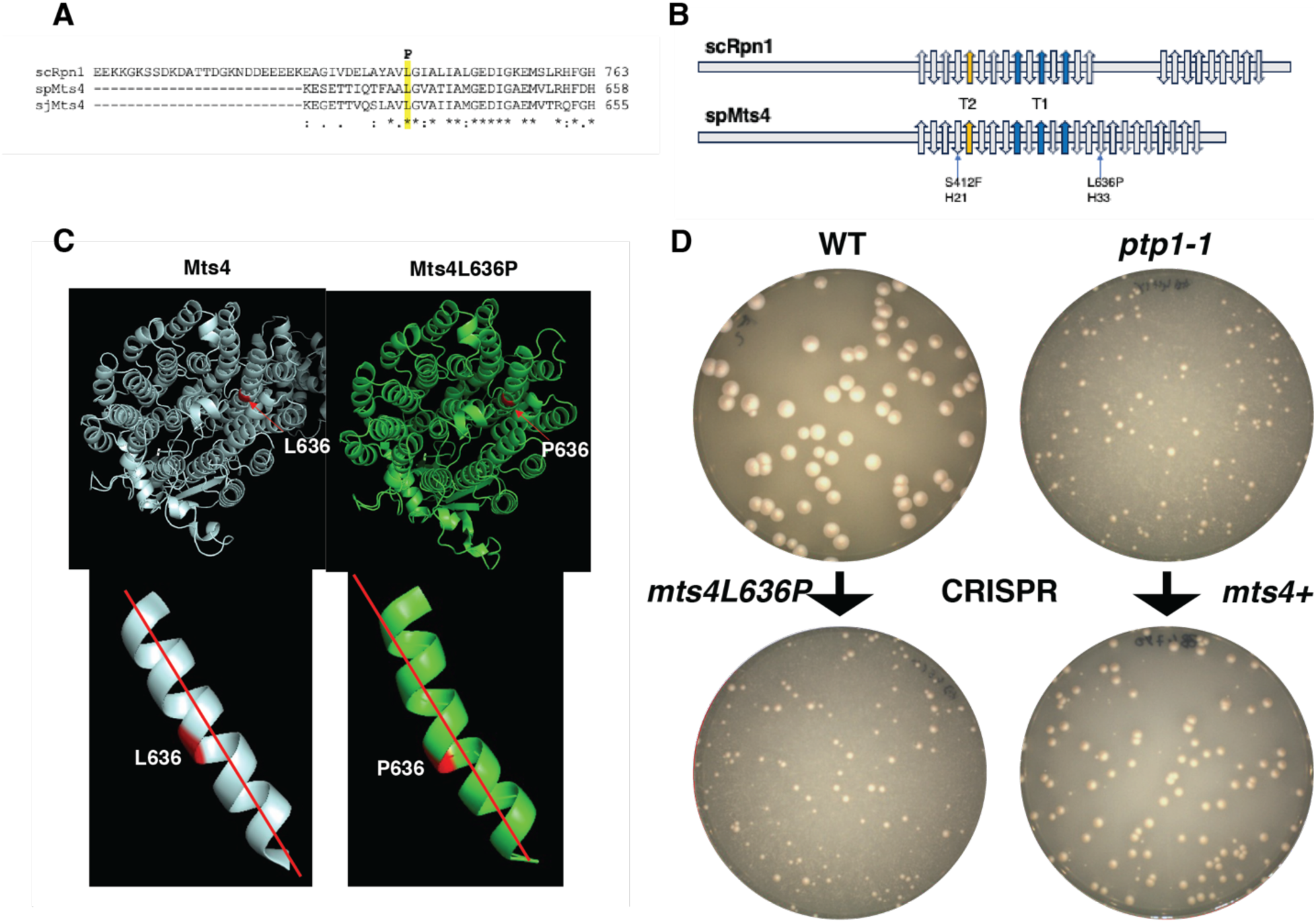
The ptp1-1 mutation is *mts4L636P*. A. Alignment of S pombe (spMts4), S cerevisiae (scRpn1) and S japonicus (sjMts4) in the region of the mutation. B. Secondary structure of scRpn1and spMts4 highlighting the positions of Mts4L636P and the Mts4S412F (Li, et al. 2019) mutations. C. Alphafold predicted structure of the Mts4L636P mutant. D. ρ0 growth assay after CRISPR mutagenesis to revert and induce the *mts4L636P* mutation.

In order to confirm that the *mts4L636P* mutation is the causative *ptp1-1* mutation, we undertook a CRISPR editing approach. We performed editing of the *mts4L636P* mutation to revert it to WT sequence in one of the *ptp1-1*/ρ+ segregants obtained in the backcrosses. In parallel, we edited the *mts4+* gene in an unrelated strain to introduce the *mts4L636P* mutation. We then tested the parental strains and the edited strains for *petite* positivity (Figure 1D). The *ptp1-1* strain reverted to *mts4+* sequence lost *petite*-positivity, but the WT strain mutated to *mts4L636P* readily yielded ρ0 cells upon treatment with EtBr. These results conclusively identify *mts4L636P* as the causative *ptp1-1* mutation.

Mts4L636P is a loss of function mutation affecting the 19S proteasome subunit The *mts4* gene codes for a central component of the 19S regulatory subunit of the proteasome. It is an essential and conserved protein, known as Rpn1 in *S cerevisiae* and PSMD2 in humans (Wilkinson, et al. 1999). L636 is located in an alpha-helical region in the domain of 11 helix-turn-helix repeats that form two concentric toroids (Figure 1A,B,C). A change to proline of a residue involved in an alpha helix is predicted to disrupt the secondary structure because the cyclic bond between the side chain and the Nitrogen in the backbone can’t form hydrogen bonds with residues in the next turn, which likely causes a significant structural change to the protein. Alphafold (Jumper, et al. 2021) modeling of the WT and *ptp1-1* mutated Mts4 protein predicts a kink in the alpha-helix where the *ptp1-1* mutation is located (Figure 1C). It is therefore likely that the Mts4L636P mutation causes a loss of function for this protein by challenging its structural stability.

The original description of the *ptp1-1* mutation left open the question of whether it represented a gain or a loss of function mutation because diploids could not be maintained in EtBr long-term cultures (Haffter and Fox 1992). The *mts4L636P* mutants grow normally as ρ+ cells. Since proteasome function is essential, this indicates that the mutation is either a mild hypomorph or a gain of function mutant. We set out to investigate the molecular phenotype of the *mts4L636P* mutation to answer this question.

We directly measured the activity of the proteasome in total cellular extracts from WT and *mts4L636P* mutants with the fluorescent chymotrypsin-like protease substrate Suc-LLVY-AMC (Figure 2A,B, C). These experiments revealed a consistent and significant decrease in MG132-sensitive peptidase activity of between 30-40% in the *mts4L636P* mutant extracts, both in rich media (Figure 2A) and in EtBr media (Figure 2C). However, when ATP was omitted from the extraction and assay buffers no difference could be detected (Figure 2B). These results indicate that the activity of the 26S proteasome, which requires ATP to maintain the interaction of the 19S and 20S subunits (Finley, et al. 2012), is compromised in the mutant, but the activity of the 20S subunit is not.

**Figure 2.**
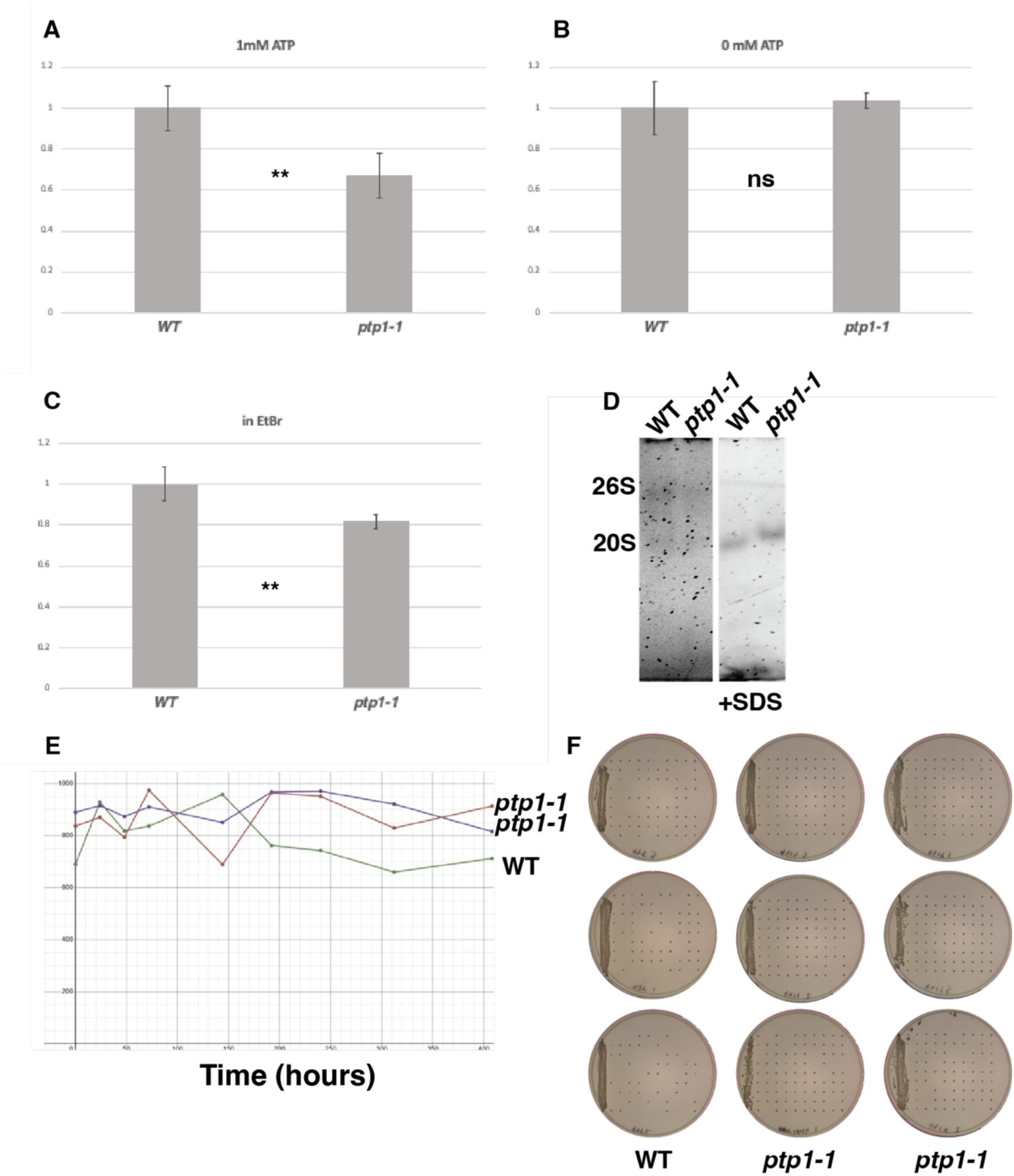
Mts4L636P is a partial loss of function allele. A, B and C: Proteasome activity assays in crude extracts with Suc-LLVY-AMC. MG132 sensitive activity is normalized to WT. A and B: Cells grown in Rich media, extracted and assayed with (A) and Without (B) 1mM ATP. C: Cells grown in EtBr media, extracted and assayed with 1mM ATP. D: In-gel activity after separation by native PAGE. E: Colony counts on 1e3 cells plated in rich media after growth in EMM media with no Nitrogen. F: Recovery of single cells dissected onto rich media after the final timepoint of growth in media with no nitrogen.

To assess the integrity of the 26S proteasome we performed an in-gel activity assay after separation of the 26S and 20S subunits by native polyacrylamide gel electrophoresis (Figure 2D). This assay showed a decreased signal of the 26S proteasome in the *mts4L636P* mutant. Together, these results show that the *mts4L636P* mutation partially compromises the activity of the 19S regulatory subunit of the proteasome.

A temperature-sensitive allele of the *mts3* component of the 19S subunit, confirmed to lose function at the restrictive temperature (Wilkinson, et al. 1999), exhibits a dramatic and rapid loss of viability when shifted to media without nitrogen that leads *S pombe* to transition to G0 stage (Takeda, et al. 2010). To test the viability of *mts4L636P* mutants in G0 we shifted it after growth to mid-logarithmic phase in rich media to media without nitrogen and maintained it in this media checking for viability by spreading on rich media at regular intervals (Figure 2E,F). Surprisingly, the viability of the *mts4L636P* mutants did not decrease even after prolonged culture and exceeded the viability of the WT after very long exposure to media without Nitrogen.

### Proteomic survey of the *mts4L636P* mutant

To assess the consequences of the partial loss of function of the 26S proteasome in cells challenged for mitochondrial function, we performed whole-proteome characterization of WT and mutant cells grown in media with EtBr. In these conditions, mtDNA replication and transcription are compromised, and growth becomes arrested. The LC/MS results detected peptides from over 4000 proteins, providing a wide survey of protein levels in these conditions (Figure 3 and Table 3).

**Figure 3.**
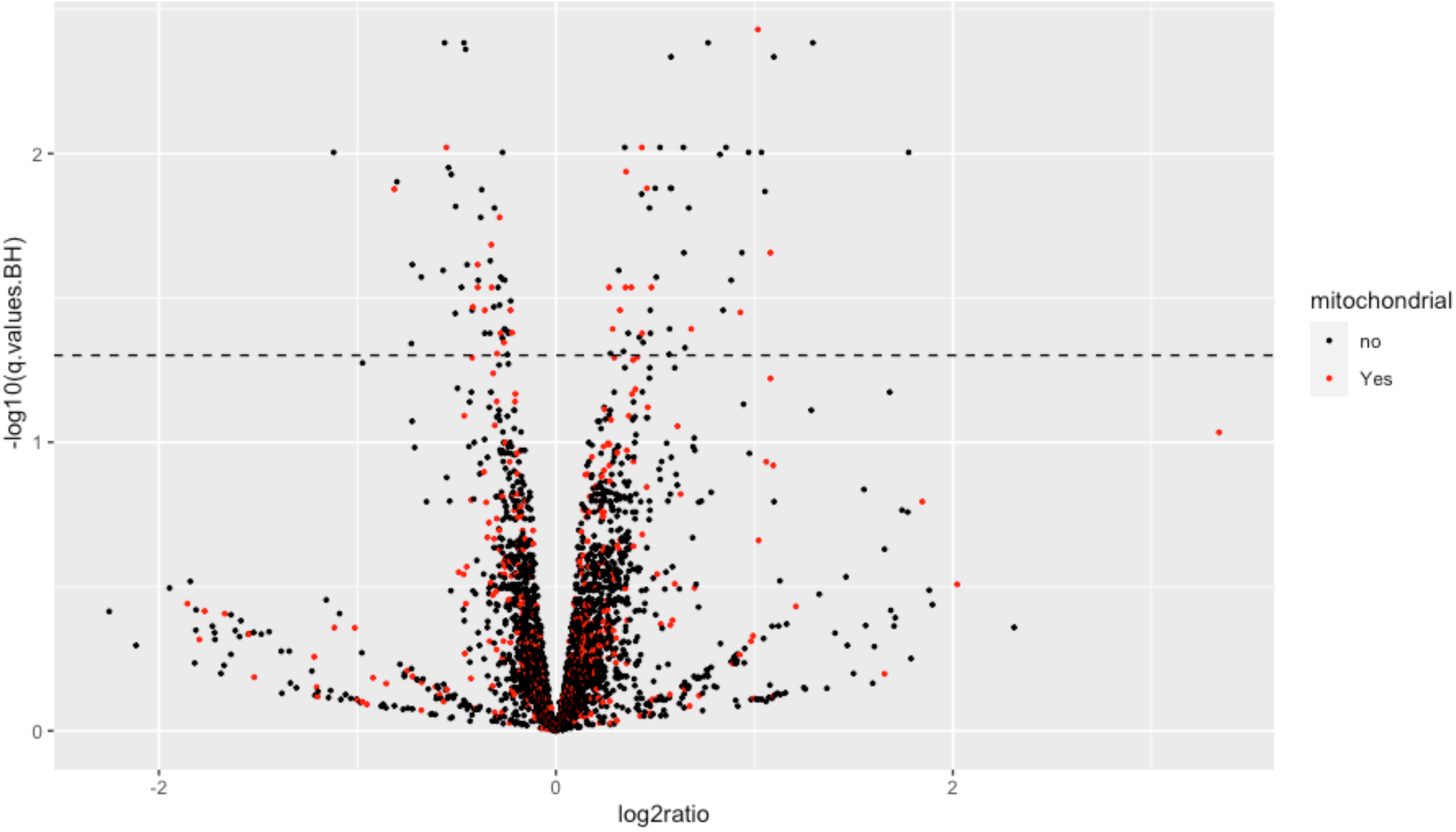
Volcano plot of Proteomics data of cells grown in EtBr media, with Log2 fold change of *mts4L636P*/WT. Mitochondrial proteins are depicted in red, non-mitochondrial in black.

**Table 3.**
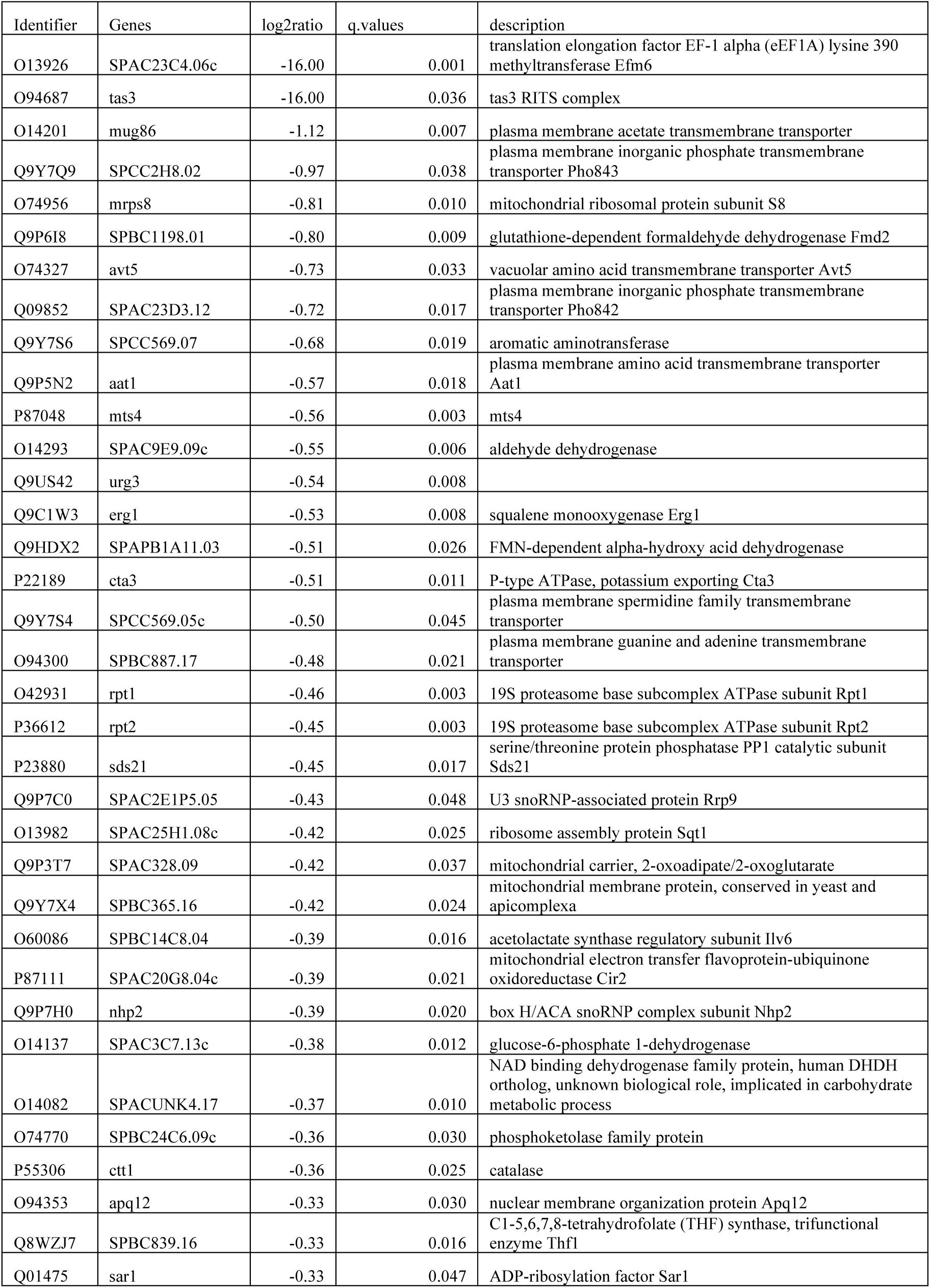
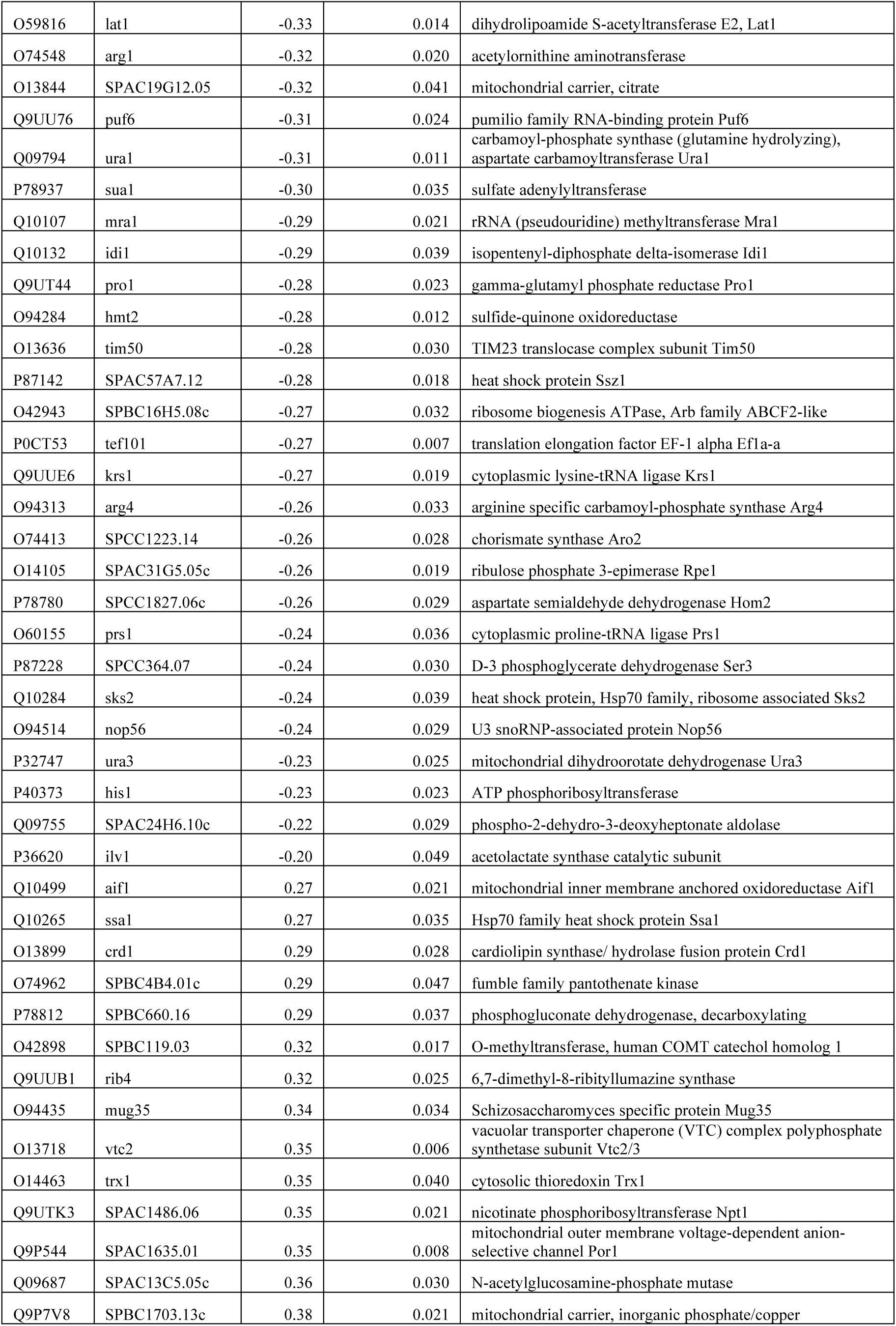
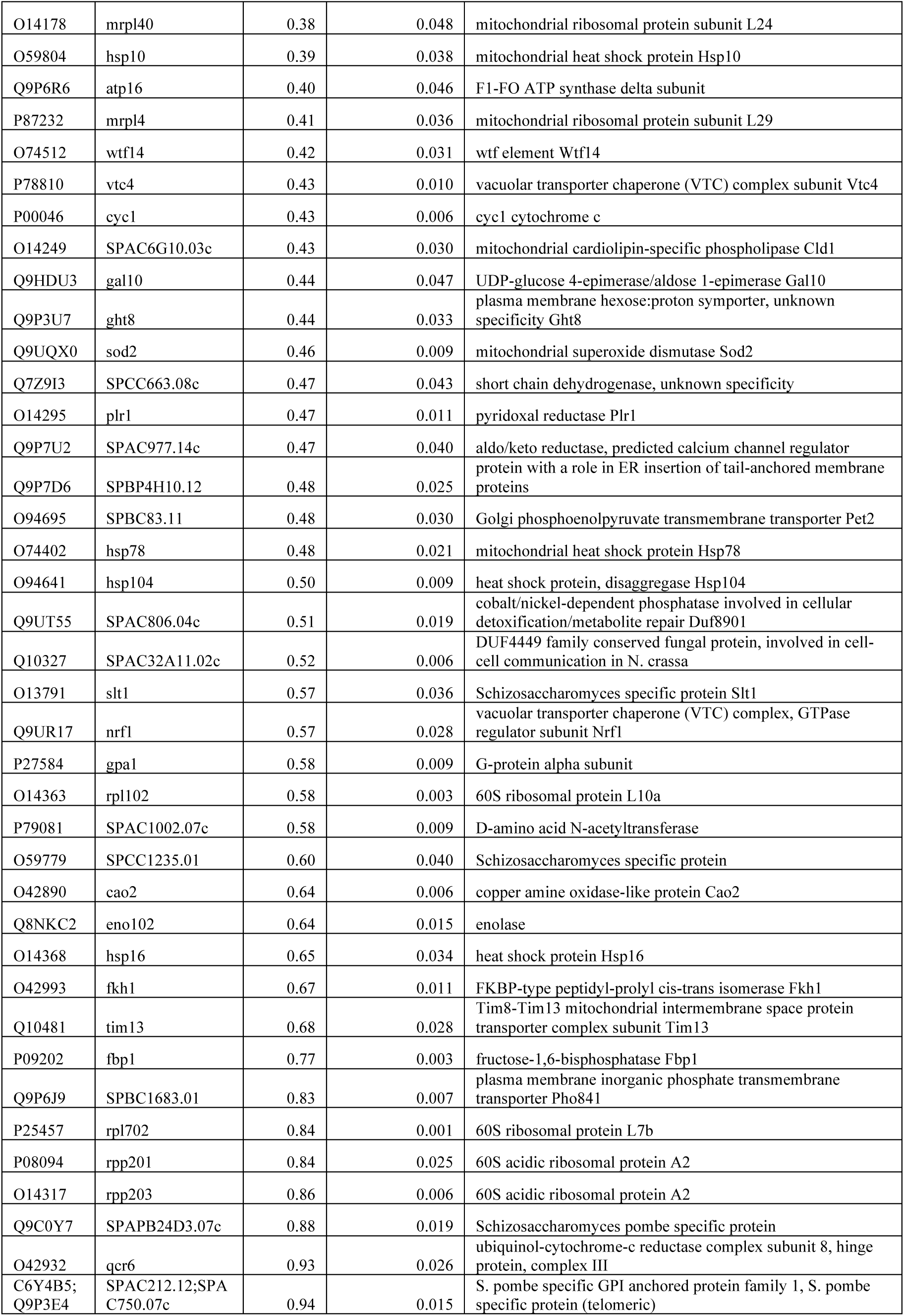
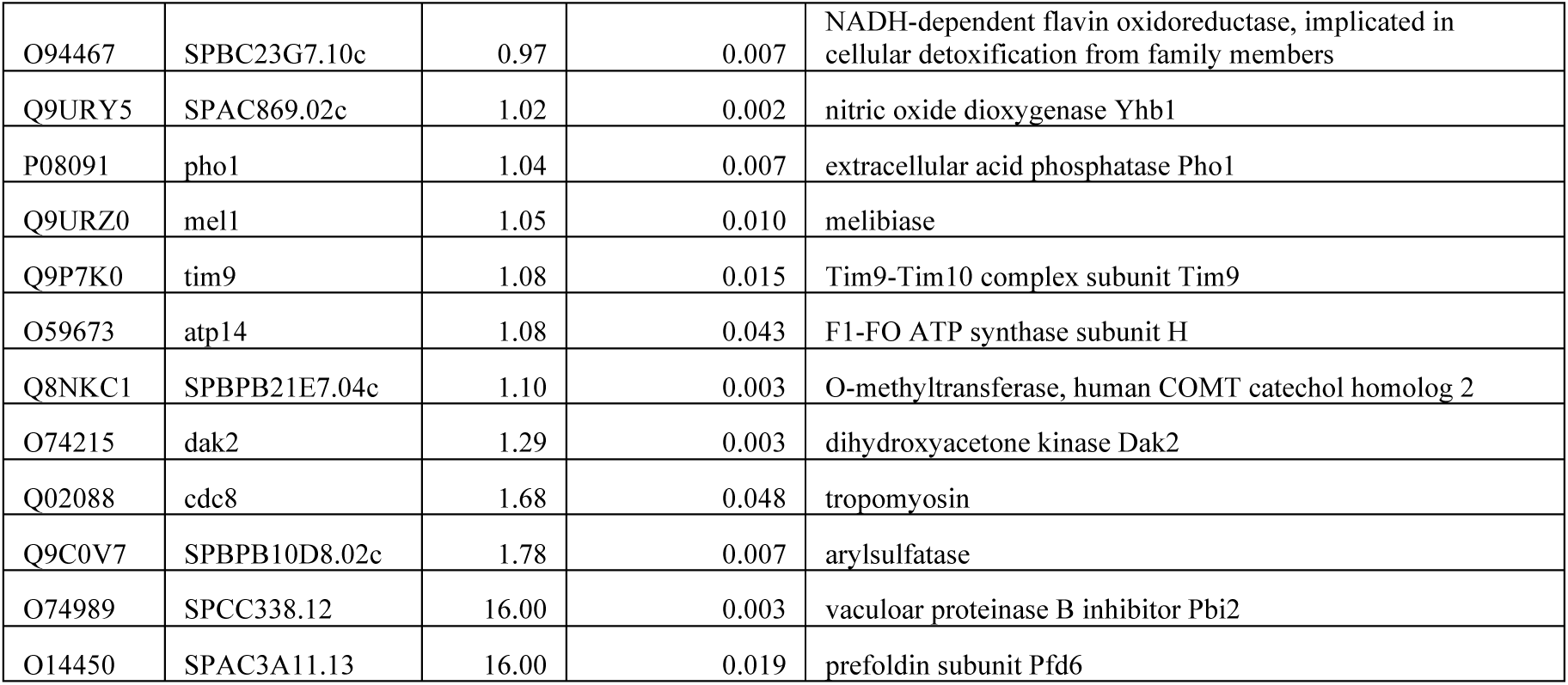
Proteomics results.

The results showed a weak effect of the *mts4L636P* mutation on protein abundances: 126 proteins exhibited significant changes. Most changes were quantitatively mild, and only 14 proteins showed differences larger than 2-fold (3 decreased and 11 increased in the *mts4L636P* mutant) (Figure 3 and Table 3). The identity of the altered proteins did not show significant enrichment in any Gene Ontologies. The proteins regulated by Ubiquitination and degraded by the proteasome would be expected to accumulate to higher levels in the *mts4L636P* mutant. Several groups of factors exhibit this behavior. While not all the proteins with altered abundance could be related to the *petite*-positivity phenotype, some groups of factors do paint compelling scenarios, and this information could be used to inform hypotheses going forward.

Of note, the abundance of Mts4, together with that of the two other 19S components Rpt1 and Rpt2, is decreased in the *mts4L636P* mutant (Table 3). These results confirm that the mutation challenges the stability of Mts4 and explain the decreased activity of the 26S proteasome.

### The *mts4L636P* mutant shows an altered oxidative stress response

Several Reactive Oxygen Species (ROS) detoxification factors are significantly down- and upregulated in the mutant, indicating an altered oxidative stress response. The mitochondrial superoxide dismutase Sod2, Thioredoxin Trx1 and the Nitric Oxide dioxygenase Yhb1 are upregulated, but Catalase Ctt1 and the Nitic Oxide detoxifier Fmd2 are downregulated (Table 3). The disfunction of the ETC upon treatment by EtBr is expected to generate superoxide ions and Hydrogen Peroxide by electron leakage from the redox centers. Sod2 detoxifies superoxide ions by partition into molecular oxygen and hydrogen peroxide. While Catalase is downregulated, the hydrogen peroxide could be detoxified by Thioredoxin and reduced Glutathione (GSH) into Oxidized Glutathione (GSSG) and water.

In this scenario, an accumulation of GSSG would be expected in the mutant. We performed a metabolomic analysis of the WT and *mts4L636P* mutant grown in EtBr media to evaluate GSH and GSSH levels. The results indicate that the mutant shows decreased levels of GSH and elevated levels of GSSG, as predicted by the model (Figure 4).

**Figure 4.**
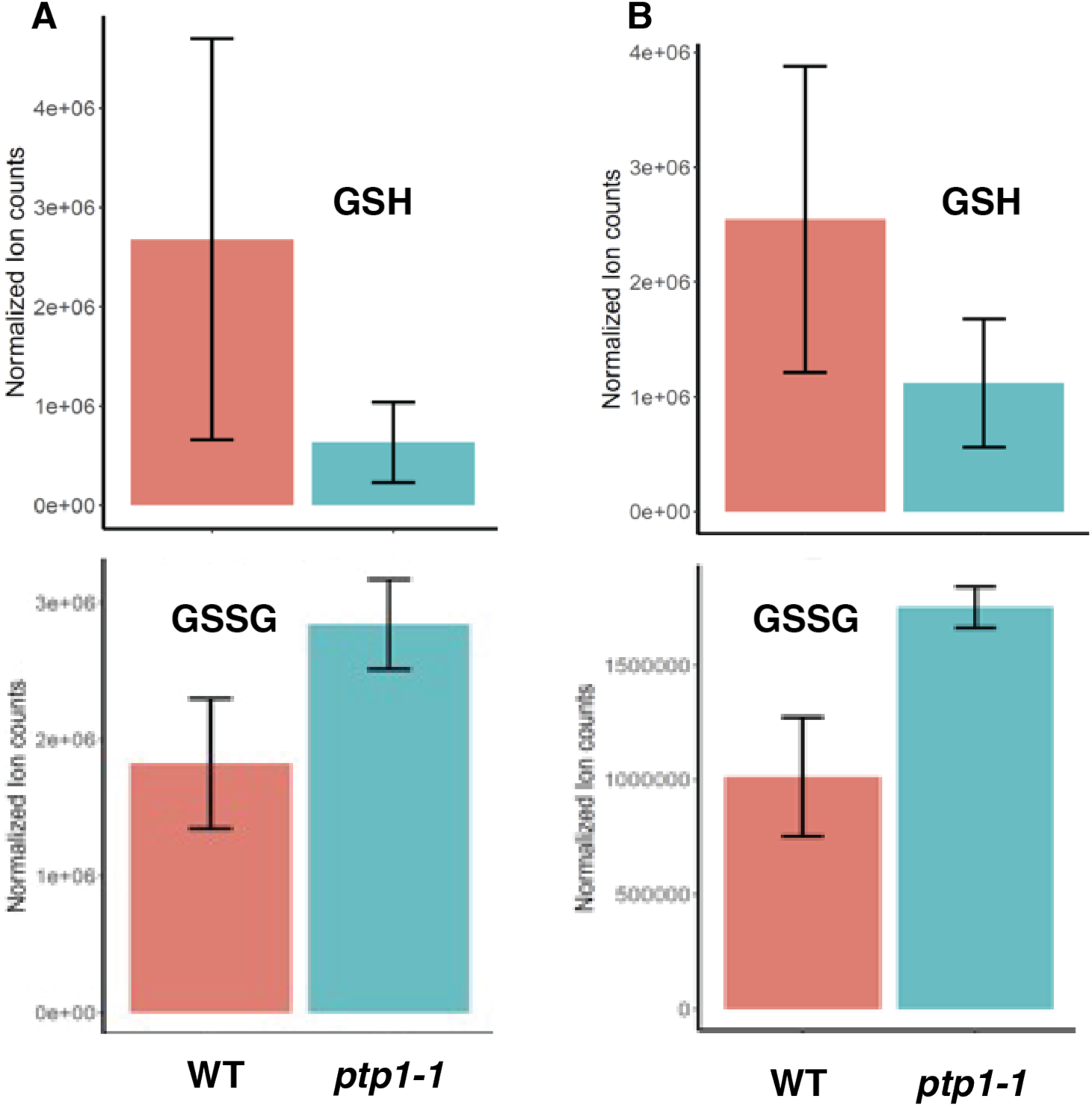
the *mts4L636P* mutation shows increased oxidized Glutathione. GSH (upper panel) and GSSH (lower panel) levels detected by mass spectrometry on negative (A) and positive (B) polarization.

### Mitophagy is not required for *petite* positivity

The temperature sensitive *mts3-1* also exhibits increased oxidative stress upon G0 arrest (Takeda, et al. 2010). This is accompanied by an increase in mitophagy that degrades damaged mitochondria. While the dramatic decrease in abundance of mitochondrial proteins observable in the *mts3-1* mutant is not replicated in the *mts4L636P* mutant, we tested whether the mitophagy pathway is involved in *petite*-positivity (Luo, et al. 2013). The mitochondrial outer membrane factor Atg43 is required for mitophagy by recruitment of Atg8 (Fukuda, et al. 2020). We crossed the *mts4L636P* mutation with the *atg43-1* allele, which abrogates mitophagy, and tested ρ0 cell generation by treatment with EtBr. *mts4L636P/atg43-1* double mutants generated ρ0 cells to levels similar to those of the *mts4L636P* single mutant, and the *atg43-1* single mutant was *petite* negative (Figure 5). These results indicate that the process of mitophagy is not involved in *petite* positivity.

**Figure 5.**
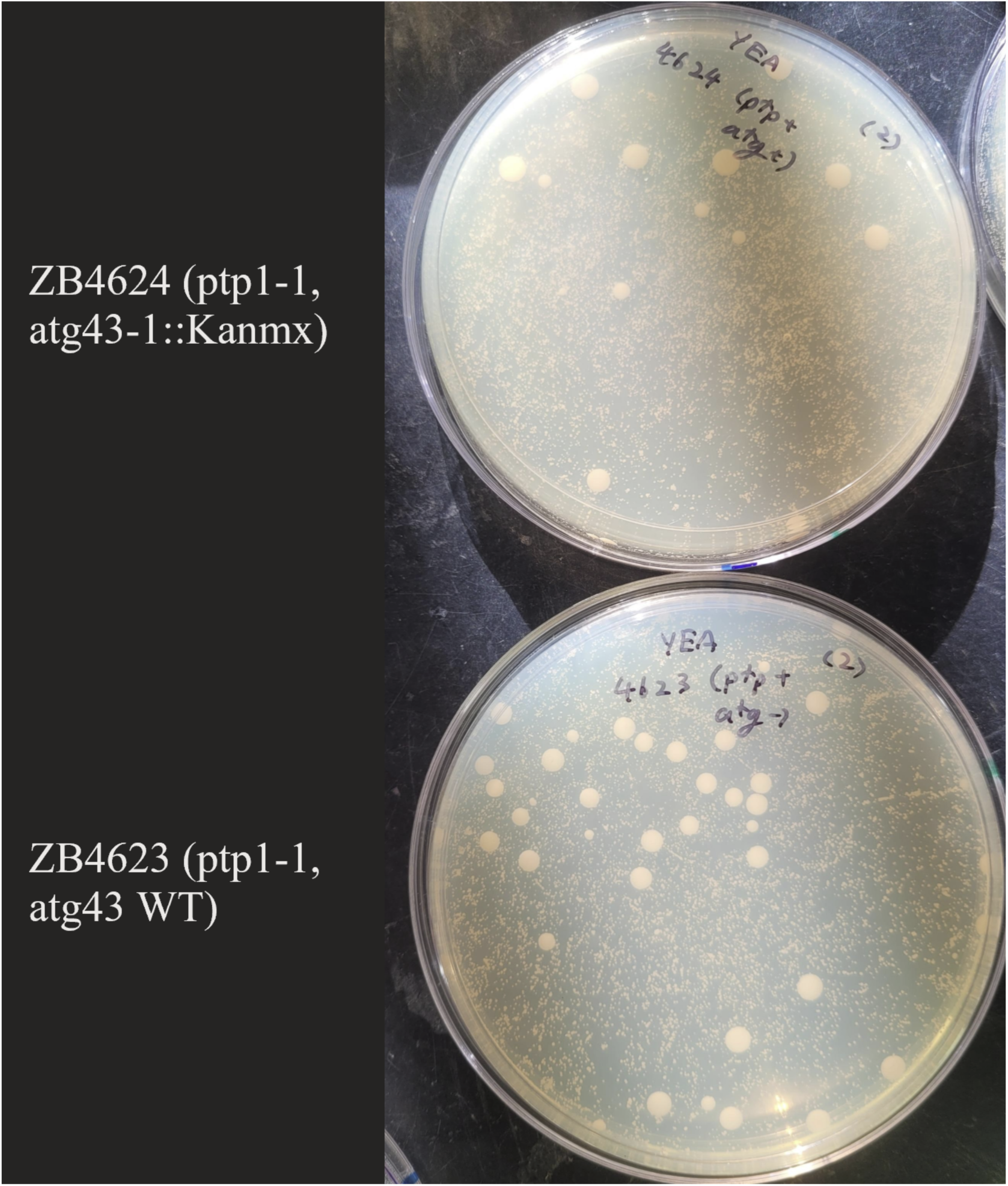
ρ0 growth assay on *mts4L636P* and *mts4L636P*/*atg43-1* strains.

## Discussion

After 75 years since the original description of *petite* mutants, the mechanisms governing survival in the absence of mtDNA are still unclear. The current consensus posits that ρ0 cells can support mitochondrial function through maintenance of ΔΨ without a functional ETC (Chen and Clark-Walker 2000; Dinh and Bonnefoy 2023). This outcome can be achieved by some species naturally, and by others through loss of regulatory pathways resulting in the increase of F_1_-ATPase ATP hydrolyzation activity (Clark-Walker 2007).

In this work, we show that fission yeast can achieve *petite*-positivity through partial loss of 26S proteasome activity. A previous large scale genetic study, focusing on the suppression of gene essentiality, uncovered that interventions that affect the proteasome, through mutation of *mts4* or overexpression of multiple proteasome-related factors, could bypass the lethality of defects in mitochondrial translation and support the survival of ρ0 cells (Li, et al. 2019). Our results confirm and extend this observation.

The Mts4L636P mutation is located in helix 33 of the toroid that binds Ubiquitinated substrates (Figure 1B), and being a L to P mutation likely disrupts the local alpha-helical structure through loss of the hydrogen bond of the amide. Li et al isolated another mutation in *mts4*, Mts4S412F, located in helix 4 of the toroid (Figure 1B) (Li, et al. 2019), that also provides *petite*-positivity. Neither of these mutations directly affect the two Ubiquitin binding sites identified in *S cerevisiae* Rpn1 (Boughton, et al. 2021), but both are predicted to dramatically affect the structure of the protein. These mutations likely destabilize the protein though misfolding. In this respect, it is interesting to note that the *mts4L636P* mutation resulted in a significant decrease in the abundance of Mts4 together with Rpt1 and Rpt2. in *S cerevisiae* these three components of the 19S proteasome subunit are assembled together as a module during proteasome maturation by the chaperone Hsm3 (Kaneko, et al. 2009; Saeki, et al. 2009). The concurrent decrease in abundance of all components of this module indicates that degradation of Mts2/Rpt1/Rpt2 likely occurs at the step of proteasome assembly.

Proteasome activity assays and native gel characterization of the proteasome isoforms indicate that the *mts4L636P* mutation affects the abundance of the mature 26S proteasome, resulting in partially decreased function. The main function of the 26S proteasome is the degradation of Ubiquitinated substrates (Finley, et al. 2012). Loss of 26S proteasome through disfunction of the 19S subunit would result in accumulation of multiple factors regulated through this pathway. It is possible that *petite*-positivity is achieved by accumulation of one or multiple protective factors normally degraded by the proteasome. Proteomic characterization of the mutant in the presence of EtBr reveals multiple candidates for such a model.

A comparison of the proteomic profiles in *mts4L636P* during growth in EtBr with that of the *mts3-1* temperature sensitive allele upon switch to nitrogen-free media (Takeda, et al. 2010) indicates that the partial loss of activity in *mts4L636P* exhibits a much milder effect on the proteome, consistent with the partial loss of function in the latter mutant. The dramatic decrease of mitochondrial protein abundance observed in *mts3-1*, owing to increased mitophagy, is not observable in *mts4L636P*. Consistently, we observed that mitophagy is not necessary for *petite*-positivity. Another difference between the effect of these two alleles is that while *mts3-1* rapidly loses viability upon a switch to G0, *mts4L636P* exhibits increased viability over WT after prolonged growth in nitrogen-free media. This might be due to exhaustion of glucose from the media, which could result in increased respiration and mitochondrial stress, revealing the beneficial effects of decreased proteasome function in these conditions.

One aspect where both *mts3-1* and *mts4L636P* coincide is the altered oxidative stress response, albeit in different ways. *mts3-1* and *mts4L636P* exhibit upregulation of multiple detoxification enzymes, with the NO dioxygenase Yhb1 present in both cases although at very different levels (Takeda, et al. 2010; Astuti, et al. 2016). But Glutathione accumulates to high levels in *mts3-1*, whereas *mts4L636P* exhibits a shift from reduced to oxidized glutathione. This could be the result of Thioredoxin Trx1-mediated detoxification of Hydrogen Peroxide, generated though higher levels of mitochondrial superoxide dismutase Sod2 and decreased levels of Catalase Ctt1. It is difficult to imagine how this altered oxidative stress response could be beneficial to the growth of ρ0 cells, or how it comes about. The proteasome directly regulates oxidative stress responses by controlling the stability of the transcription factor Pap1 (Marte, et al. 2020), but this protein was not elevated in the proteomic profile of the *mts4L636P* mutant. Interestingly, oxidative stress regulates proteasome function by direct glutathionylation of the 20S subunit (Demasi, et al. 2014), which becomes more efficient in its function to degrade oxidized and misfolded proteins independently of ATP and the 19S subunit (Grune, et al. 2003). Since the extreme mitochondrial stress resulting from loss of the mtDNA would decrease cellular ATP levels, increase ROS, and result in the cytoplasmic accumulation of mitochondrial protein precursors, such a scenario could provide a possible mechanism for *petite* positivity.

Another set of factors that show upregulation in both *mts3-1* and *mts4L636P*, providing possible candidates for Ubiquitination-regulated targets, are cytoplasmic and mitochondrial chaperones. Some of these might become indirectly upregulated as part of the Unfolded Protein Response pathway due to the mitochondrial stress present in both conditions (Narayana Rao, et al. 2022), which could be exacerbated by proteasome disfunction, but they provide a list of possible direct targets of the proteasome as candidate mediators of *petite* positivity. Of particular interest are the mitochondrial matrix chaperones Hsp78 and Hsp10, and the intermembrane chaperones Tim9/Tim10 and Tim13, which assist protein import into the inner membrane and the matrix and their correct folding after reaching their destination. The proteasome regulates the abundance of intermembrane proteins prior to their import (Bragoszewski, et al. 2013; Kramer, et al. 2021) and the structure of Tim9 and Tim10 is regulated by the redox state of the cell through the formation of cysteine disulfide bridges (Morgan and Lu 2008). Furthermore, Tim9/10 are known to be regulated by Yme1 (Spiller, et al. 2015), an known mediator of *petite*-positivity (Thorsness, et al. 1993; Kominsky and Thorsness 2000). If the defective mitochondrial protein import in ρ0 cells can be corrected by increased activity of mitochondrial chaperones, it would be beneficial to their survival. Other mitochondrial factors accumulating in *mts4L636P* could also explain the survival after loss on mtDNA in a manner analogous to that observed in mutants of the F_1_-ATPse subunits. In particular, the F_1_-ATPase components Atp14 and Atp16 (subunits H and δ, respectively) are moderately upregulated, but the function of these subunits is not clear and their influence on ATP hydrolysis by the F_1_ subunit is hard to predict (Duvezin-Caubet, et al. 2006). The wide involvement of the proteasome in virtually every cellular process makes it difficult to pinpoint a mechanism by which its decreased activity would ameliorate mitochondrial function. This study sets the stage for further investigation of this phenomenon, which could have profound implications in our understanding of mitochondrial biology and its consequences on aging.

## Author contributions

KLA, LH, SC and MZ performed experiments. KLA and MZ analyzed data and wrote the manuscript.

## Acknowledgements

We are grateful to Tom Fox, Li-lin Du and Nathalie Bonnefoy for reagents and strains, and to Kiran Madura and Steven Brill for technical advice. Research reported in this manuscript was supported by the National Institute of General Medical Sciences of the National Institutes of Health (NIGMS/NIH) under grant number R35GM131763. This work has not undergone peer review.

## Competing interests statement

The authors declare no competing interests.

## Notes

### Competing Interest Statement

The authors have declared no competing interest.

